# Neopolyploidy increases stress tolerance and reduces fitness plasticity across multiple urban pollutants: support for the ‘general purpose’ genotype hypothesis

**DOI:** 10.1101/2023.09.08.556614

**Authors:** Martin M. Turcotte, Nancy Kaufmann, Katie L. Wagner, Taylor A. Zallek, Tia-Lynn Ashman

**Affiliations:** Department of Biological Sciences, University of Pittsburgh, Pittsburgh PA 15260

**Keywords:** adaptation, urban ecology, urban evolution, environmental pollutants, Lemnaceae, stress tolerance, environmental trade-off, plasticity in fitness, genetic correlation, autopolyploidy

## Abstract

Whole genome duplication is a common macromutation with extensive impacts from gene expression, to cellular function, and whole organism phenotype. As a result, it has been proposed that polyploids have ‘general purpose’ genotypes that perform better than their diploid progenitors under stressful conditions. Here we test this hypothesis in the context of stresses presented by anthropogenic pollutants. Specifically, we tested how multiple neotetraploid genetic lineages of the Greater Duckweed (*Spirodela polyrhiza*) perform across a favorable control environment and five urban pollutants (iron, salt, manganese, copper, and aluminum). By quantifying the population growth rate of duckweed over multiple generations we found that across most pollutants, but not all, polyploidy decreased the growth rate of actively growing propagules but increased that of dormant ones. Yet, when considering total propagule production, polyploidy increased tolerance to most pollutants and polyploids maintained population-level fitness across pollutants better than diploids. Furthermore, broad-sense genetic correlations in growth rate among pollutants were all positive in neopolyploids but not so for diploids. Our results provide a rare test and support for the hypotheses that polyploids are more tolerant of stressful conditions and can maintain fitness better than diploids across heterogenous stresses. These results may help predict the distribution of polyploids across stress gradients such as those caused by urbanization and other human activities.

## INTRODUCTION

Polyploidy (the possession of more than two sets of each chromosome resulting from whole genome duplication) is a major macromutation that occurs throughout eukaryotes, especially in plants where most have at least one genome duplication in their past (e.g., Wood *et al*., 2009). Polyploids often show increased genetic diversity, genomic plasticity and phenotypic novelty relative to their diploid progenitors (Leitch & Leitch, 2008; Soltis *et al*., 2009; Van de Peer *et al*., 2020) allowing them to rapidly adapt and/or invade new habitats (e.g., te Beest *et al*., 2012; Pyšek *et al*., 2023). Interestingly, polyploidy is often associated with historical periods of environmental change (Fawcett *et al*., 2009; Van de Peer *et al*., 2017) as well as with harsh or stressful habitats globally (Ehrendorfer, 1980; Rice *et al*., 2019). Indeed, it has long been proposed that polyploids might have ‘general purpose’ genotypes that are beneficial across a variety of circumstances (Baker, 1965; Stebbins, 1971). Recently these ideas have been combined to suggest that polyploids may perform better than diploids in the face of contemporary anthropogenic stressors, such as those found in urban environments (Van Drunen & Johnson, 2022).

Urban environments, with their vast infrastructure, impervious surfaces, and human and industrial pollutants represent a confluence of anthropogenic forces. Plants in urban habitats often experience increased abiotic stresses such as elevated temperature, altered water and nitrogen availability, novel chemical pesticides and industrial waste (Parris, 2016; Van Drunen & Johnson, 2022). Contaminants that can be detrimental to plant growth such as alkaloids (e.g., aluminum), heavy metals (copper, iron and manganese) and salts (NaCl), among many others, enter soils and waterways through un- or partially treated effluent, atmospheric deposition, and stormwater runoff from impervious surfaces (Hope *et al*., 2004; Jacobson, 2011; Ruas *et al*., 2022). While some contaminants are micronutrients (e.g., Mn, Fe) that play essential roles in the plant life cycle, excesses and unbalances can induce cytotoxic and genotoxic effects and reduce plant fitness (Dutta *et al*., 2018). Because polyploids can be more tolerant than diploids to a wide range of abiotic stresses, e.g., thermal, drought, saline stress, nutrient deficiencies and excesses (Van de Peer *et al*., 2020; Tossi *et al*., 2022) they may have growth advantages in these contaminated environments. Surprisingly, pollution and micronutrient excess are the least studied urban stresses to plants (Ruas *et al*., 2022) and also least studied from the perspective of ploidal tolerance differences (Tossi *et al*., 2022). Thus, an important step towards testing the hypothesis that polyploids will have an advantage in urban habitats via increased stress tolerance (Van Drunen & Johnson, 2022) will be to compare diploid and polyploid fitness under a suite of common urban contaminants.

Stress tolerance is achieved via a variety of physiological, molecular and morphological mechanisms (reviewed in Tossi *et al*., 2022) which may impact whether ploidy alters tolerance of different stressors in a specific or general manner. For instance, multiple detoxifying and sequestration mechanisms exist in plants (Dutta *et al*., 2018). And although polyploids were found to have greater tolerance in 90% of reviewed studies (Tossi *et al*., 2022), the effect of polyploidy can at times lead to lower stress tolerance. For example, Mouhaya *et al*. (2010) found increased sensitivity to saline stress in natural autotetraploid citrus species. Moreover, since few studies test the independent effects of multiple stressors (but see Bafort *et al*., 2023; Mattingly & Hovick, 2023), the question of whether polyploidy provides general stress tolerance in any given system is not yet been answered. In principle, however, because increased gene copies, allelic diversity or network flexibility can allow polyploids to produce a greater quantity or variety of gene product (Parisod *et al*., 2010; De Smet & Van de Peer, 2012) one would predict that polyploidy would generally increase tolerance to a variety of stressors, especially where greater enzymatic activity increases detoxification or increased transmembrane transport upregulates sequestration. Additionally, larger cell or vacuole size of polyploids (Doyle & Coate, 2019) may allow for greater storage of toxic substances as seen in hyperaccumulator species (Leitenmaier & Küpper, 2011). Moreover, when mechanisms of tolerance are the same for two stressors (e.g., both involving the same antioxidant pathway; Dutta *et al*., 2018) polyploidy may have a similar effect on tolerance to each. However, if there are tradeoffs between tolerance mechanisms in diploids, for example if the mechanisms to cope with toxicity from two stressors are alternate pathways, or rely on the same resource pool, we may find that the greater genomic plasticity or genetic variation afforded by a polyploid genome (Leitch & Leitch, 2008) buffers against these tradeoffs, leading to less negative or even overall positive correlation among stress responses for polyploids. Alternatively, if the underlying mechanisms (e.g., uptake limitation versus antioxidative to detoxify) or genetic determinants (e.g., allelic diversity or gene interactions) vary among stressors then we might find no universal pattern in the effect of polyploidy on stress tolerance. Thus, it is still an open question as to whether polyploids have ‘general purpose’ genotypes (Baker, 1965; Stebbins, 1971) that buffer them against a wide range of stressors, including those commonly encountered in urban settings.

The ultimate expression of stress tolerance is maintaining fitness in the face of a particular stressor (Simms, 2000). A polypoid possessing a general purpose genotype is predicted to exhibit fitness homeostasis (e.g. lack of fitness plasticity) across multiple environments (Madlung, 2013; Wei *et al*., 2019; Mattingly & Hovick, 2023). However, most studies comparing stress tolerance between diploids and polyploids compare enzymatic metrics, physiological responses or fitness proxies of growth or development (e.g., reviewed in Tossi *et al*., 2022) to a single stressor and have not addressed fitness homeostasis across a range of stressors (but see Wei *et al*., 2019; Mattingly & Hovick, 2023). Because stress may impact different aspects of plant life-history these may not fully be captured by measuring only physiological responses or biomass change in individuals whereas as fitness integrates all impacts. The most appropriate fitness comparison is to be made at the population level, where diploid and polyploid growth both within generations and reproduction among generations contribute to population fitness (e.g., Selmecki *et al*., 2015; Anneberg *et al*., 2023).

Recurrent formation of polyploids from genetically different diploids is a common phenomenon in nature (Soltis & Soltis, 1999; Kolář *et al*., 2017) and could lead to variation in stress tolerance even when polyploidy involves one parental genome (autopolyploidy) rather than two genomes (allopolyploidy) (e.g., Wei *et al*., 2019; Bafort *et al*., 2023; Mattingly & Hovick, 2023). This variation in stress tolerance may ultimately determine the probability of polyploid establishment under novel or stressful conditions (Soltis & Soltis, 1999; Van Drunen & Johnson, 2022). While use of multiple sources of natural diploid and polyploids can address this to some degree (e.g., Wei *et al*., 2019), synthetic neopolyploids are recognized as a powerful tool because they avoid the confounding effects of evolution after duplication that exist in wild polyploid–diploid comparisons (Bomblies, 2020). Furthermore, including multiple genotypes of synthetic autopolyploids allows for evaluating the potential for genetic diversity to contribute to stress tolerance (co)variation in diploids and to test the effect of polyploidy on these relations.

Here we used diploids and neopolyploids of the model plant the Giant Duckweed (*Spirodela polyrhiza;* Lemnaceae). These floating aquatic angiosperms mostly reproduce clonally and have a rapid generation time of 4-5 days (Acosta *et al*., 2021) making them a proven system for experimental population-level studies (Armitage & Jones, 2019; Hart *et al*., 2019; Hitsman & Simons, 2020; Subramanian & Turcotte, 2020; Huber *et al*., 2021; Hess *et al*., 2022) and several diploid duckweed species are being studied for stress responses at the phenomenological and mechanistic levels and used as potential phytoremediators (Dalu & Ndamba, 2003; Huber *et al*., 2007; Ziegler *et al*., 2017; Chmilar & Laird, 2019; Ekperusi *et al*., 2019; O’Brien *et al*., 2022).

We grew six genetically distinct pairs of diploids and their immediate neotetraploid descendants (Anneberg *et al*., 2023) individually in water contaminated with one of five urban pollutants (copper, iron, salt, aluminum, and manganese) and a control to answer these specific questions concerning stress tolerance.

1. How does environmental pollution alter the relative fitness of diploids and their derived neotetraploids? Does it depend on genetic lineage or pollutant?
2. Are neotetraploids more stress tolerant than diploids, or does it depend on genetic lineage or pollutant?
3. Do neotetraploids maintain fitness (lower fitness plasticity) across pollutants better than their progenitor diploids?
4. Are there genetic tradeoffs in fitness across pollutants and is this altered by polyploidy?

## METHODS

### Study System and the Creation of Neopolyploids

*Spirodela polyrhiza (L.) Schleid,* is a globally distributed small floating aquatic plant in the family Lemnaceae (Jacobs, 1947). They exist in fresh water ponds, streams, as well as being common in urban parks, drainage ditches, and waste water collection sites where they encounter numerous pollutants (Dalu & Ndamba, 2003; O’Brien *et al*., 2022). When reproducing asexually through budding, they can produce actively growing individuals (hereafter referred to as fronds) or under stress produce dormant propagules termed turions (Jacobs, 1947; Appenroth *et al*., 1996).

In 2017, *S. polyrhiza* was sampled from natural and urban ponds in Western Pennsylvania and Eastern Ohio, U.S.A. (Table S1). Individuals were used to establish six mono-clonal colonies and were confirmed to be unique genotypes using 10 microsatellite loci (Table S1; Xu *et al*., 2018; Kerstetter *et al*., 2023). These were cultured in conditions that maintain asexual reproduction and formed the initial genetic lineages. Synthetic neotetraploids were created from six genetically distinct diploids using the mitotic inhibitor colchicine (Sigma Aldrich, CAS: 64-86-8). Details of the methodology are found in Anneberg *et al*. (2023), and four of the lineages used here are reported therein. Briefly, after exposing populations of a single diploid genotype to colchicine we tested ploidy using flow cytometry following Wei *et al*. (2020). We retained both converted neotetraploids and unconverted diploids from each genetic lineage. These were maintained in quarter strength growth media described in Appenroth *et al*. (1996). Before the experiment, each of the 12 sublineages (a diploid and polyploid of each genetic lineage) were grown in common garden conditions for two weeks. This consisted of growth in full strength media under fluorescent lights at room temperature.

### Experimental Design

We selected five urban pollutants that duckweed populations may commonly encounter (Bhat, 2012; Vo *et al*., 2018; O’Brien *et al*., 2022). The concentrations used during the experiment were determined by pilot studies on other duckweed diploid genotypes (Zallek and Turcotte unpublished). We selected concentrations that reduced population growth but did not kill all the duckweed within a few weeks. Given that our focus is to compare among ploidies rather than among pollutants per se we did not aim to equalize the negative impacts of each pollutant. Each experimental unit consisted of 120 mL glass jar (Fisher, U.S.A., # FB02911775) to which 90 mL of quarter strength autoclaved media was added. Jars were set in plastic trays and a large plastic lid covered the tray. Pollutant treatments varied in the addition of nothing (control), 0.04 mM of FeCl_3_ (iron), 40 mM of NaCl (salt), 0.6 mM of McCl_2_ (manganese), 0.0025 mM of CuSO_4_ (copper), or 0.015 mM of Al_2_(SO_4_)_3_ (aluminum). The growth media also contains small quantities of Fe (0.025 mM), Mn (0.013 mM), Na (0.0258 mM) and Cl (0.026 mM) as micronutrients. During the week of January 23, 2023, four individual duckweed (fronds) were added to each jar. On days 7, 14, and 21 jars were photographed and duckweed fronds were manually counted from the photos using Fiji (Schindelin *et al*., 2012). Turions were counted on day 21. We counted fronds and turions separately because the relative performance of diploids and neopolyploids can depend on which is quantified (Anneberg *et al*., 2023).

The experiment was conducted by 836 undergraduate students at the University of Pittsburgh taking a research focus lab course. Students were divided into 42 course sections of up to 20 students, taught by 13 instructors, across three laboratory classrooms. Genetic lineages were tested across different rooms and instructors, but each section tested only two lineages. Sublineages (diploid and polyploid) of a specific genetic lineage were always tested together. Specifically, students worked in pairs, setting up four jars of a single lineage (e.g., SP.05) in a factorial design of control or a single pollutant crossed with the diploid or polyploid sublineage. This approach led to an unbalanced design with five times as many control jars but was important to teach students the importance of controls. Thus, each section set up 24 control jars and four jars of each metal. After removing jars with missing data or major errors (e.g., adding greater than 50% too many duckweed initially) a total of 1591 experimental units (jars) were analyzed. Each sublineage by pollutant combination (excluding control) had on average 13.25 (SD = 3.08) replicates.

### Statistical Analysis

To address the relative performance of ploidies under varied pollutants we separately quantified the production of fronds and turions. For fronds, we first compared the fit of linear and exponential population growth models. In these models, the response variable was the abundance of fronds and explanatory fixed factors included ploidy, genetic lineage, and pollutant as well as their interactions. In all models the initial abundance was set as the initial frond number added to each jar, (i.e., not estimated by the model). Jar was a random effect that accounted for repeated measures. We tested various models with different random effects (section number, instructor, or room) as well as the presence or absence of a variance function that increased with time. The best fitting model, as determined by model comparison using AIC, was a linear model with jar nested within section number as random effects. Models were fit using the nlme function in the package of the same name (Bates & R Core Team, 2023). In addition to the general statistical analysis, which used type III sums of squares, we conducted planned contrasts comparing the growth rates of diploids versus polyploids within each pollutant using the emmeans package (Russell, 2023).

We tested for differences in turion production using a linear mixed-effect model with turion as the response variable, with fixed factors of ploidy, genetic lineage, and pollutant as well as their interactions, and section number as a random effect using the lme function in the nmle package. We then conducted planned contrasts as above.

To evaluate ploidal differences in tolerance to pollutants and fitness plasticity across pollutants we analyzed the total abundance of individuals (fronds and turions combined) to provide an overall assessment of population growth. To evaluate tolerance and fitness maintenance we first quantified the growth rate within each jar. We fit a simple linear population growth model to each jar with initial abundance pre-determined by the initial number of individuals added. Then we calculated tolerance as the growth rate in a given pollutant divided by the control, where each pair of jars was measured by a single student and represented one control and one single pollutant of the same lineage and ploidy. We fit a linear mixed-effect model to these tolerance values with ploidy, lineage, pollutant, and their interactions as fixed effects and section number as a random effect. We also conducted planned contrasts comparing diploids to polyploids across all pollutants as well as in each pollutant treatment.

To quantify fitness plasticity in the face of the five pollutants, excluding the control, we also analyzed variation in growth rate at the jar level. We calculated the Relative Distance Plasticity Index (RDPI, Valladares *et al*., 2006) for each sublineage (combination of lineage and ploidy). The relative differences in growth rate among two jars is the absolute difference in growth rates divided by the sum of both growth rates. This is calculated for all possible pairings of jars that have the same duckweed sublineage but only between jars that have a different pollutant treatment (e.g., diploid SP.05 iron and salt). Then the relative differences are summed and divided by the number of pairings to give the RDPI value. This is then repeated for each sublineage. Using these distances, we fit a linear model with lineage and ploidy and their interaction as fixed effects, we then used planned contrasts to compare ploidy within each lineage. We used scripts from Ameztegui (2017) to run our analyses.

We then calculated broad sense genetic correlations among pairs of pollutant. First, we fit a mixed-effect model to the jar level growth rate data that had ploidy, lineage, pollutant and their interactions as fixed effects and section number as a random effect. Given that the duckweed reproduce clonally, we calculated genetic correlations among each pair of pollutants using the estimated marginal means from the mixed-effect model. We calculated Pearson correlation coefficients and their significance among the six diploid lineages for each possible pair of pollutant and repeated this procedure for the polyploids. We used the cor function in the base package of R as well as the corrplot package (Wei & Simko, 2021). Finally, we used a paired t-test on the correlation coefficients to test whether the correlations were differed among diploids and polyploids.

## RESULTS

### Environmental pollution alters the relative performance of diploids and their derived neotetraploids in ways that depend on pollutant and genetic lineage

The effect of ploidy on performance quantified as both the growth rate in frond abundance as well as the total number of turions produced (see Figure S1 for time series) had pollutant- and genetic lineage-dependent effects. A linear mixed-effect model revealed that ploidy, lineage, and pollutant and all interactions significantly impacted the growth rate of fronds (all P < 0.001; Table S2; Fig. 1). Although interpretation is complex given the interactions, planned contrasts (Fig.1) revealed that the relative performance of the diploids depended both quantitatively (effect size) and qualitatively (direction of effect) on the pollutants. Neotetraploids performed significantly worse in control than diploids (−37% in population growth rate, see Fig. 1 for P values), in aluminum (−31%), iron (−30%), and in salt (−18%). Yet, in manganese performance did not differ significantly (−8%) and in copper the polyploids performed significantly better (+18%). While there was variation among genetic lineages, they generally followed these overall trends (Fig. 1). In control, iron, salt, and aluminum diploid populations grew faster than their derived polyploids for all lineages but the effect sizes varied. But in copper and manganese polluted media, approximately half the genetic lineages show polyploids performed better than diploids. Moreover, when we fit models to each pollutant separately, there were significant ploidy by lineage interactions (P < 0.001) in all stressors except for iron and aluminum (P = 0.18 and P = 0.14 respectively).

**Figure 1:**
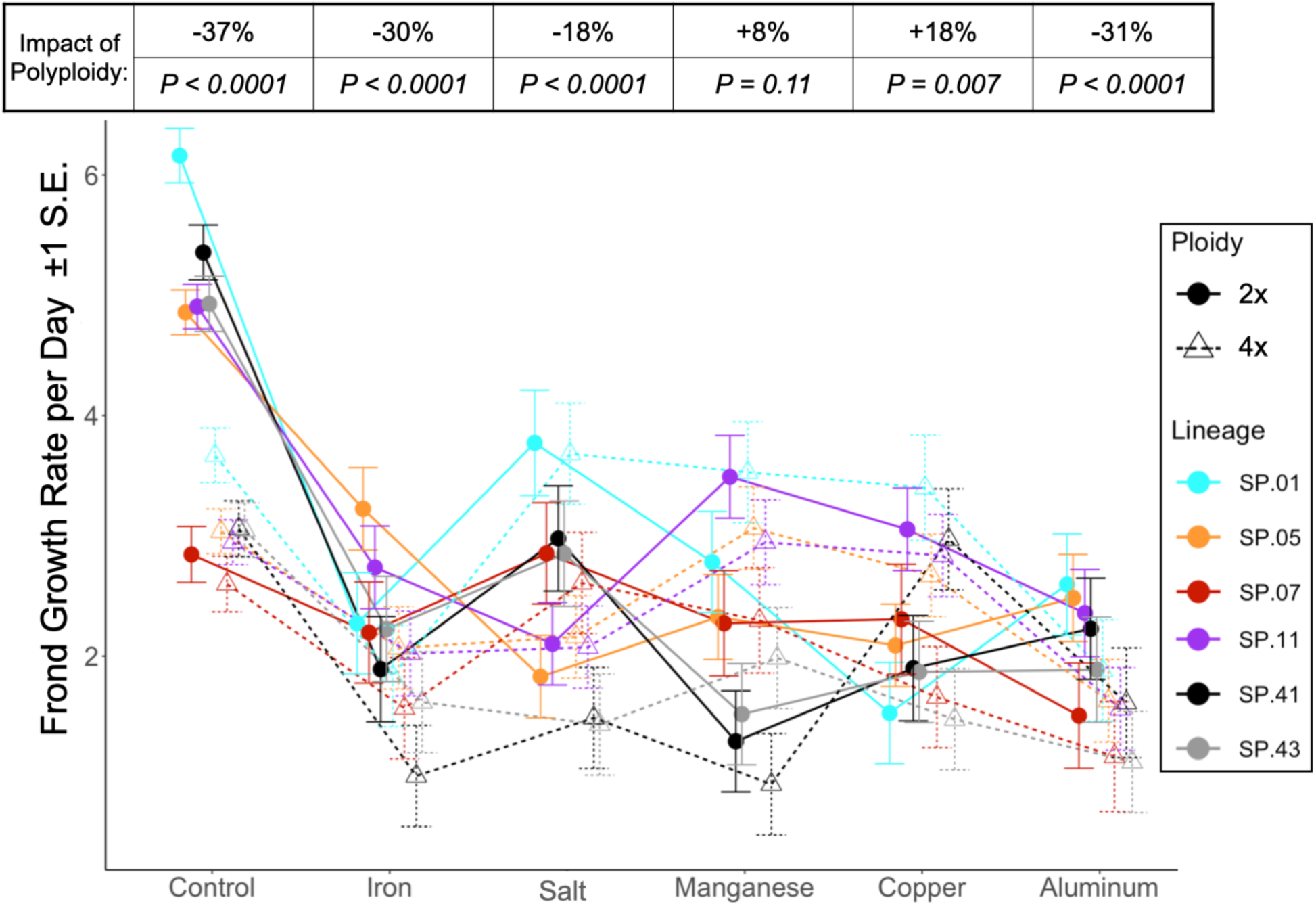
Daily frond population growth rates (estimated marginal means ± 1 S.E.) separated by ploidy and genetic lineage across the six pollutant treatments (including the control). The table above shows the % difference of neopolyploid (4x) relative to diploid (2x) and P values from the six planned contrasts between diploids and neotetraploids for each pollutant treatment. Point positions were dodged for clarity.

In contrast to fronds, turion production was higher in neopolyploids than diploids but was also strongly impacted by genetic lineage and pollutant type (Fig. 2; Fig. S1). Unlike frond growth rate, the control conditions lead to similar total turion production even though there were generally more fronds in the control treatment. The mixed model revealed strong effects of genetic lineage and its interactions with ploidy and pollutant (all P ! 0.0001; Table S3). Yet, averaging across lineages, planned contrasts show that polyploids produce more turions in iron (+196%; Fig. 2), aluminum (+131%), copper (+94%), and in control (+29%), whereas they do not differ significantly from diploids in manganese and salt. Pollutant specific models revealed all treatments had significant ploidy by lineage interactions (salt P = 0.049, all others P < 0.0001). Indeed, salt almost completely prevented turion production for both diploids (estimated marginal means of 0.012) and neopolyploids (0.139).

**Figure 2:**
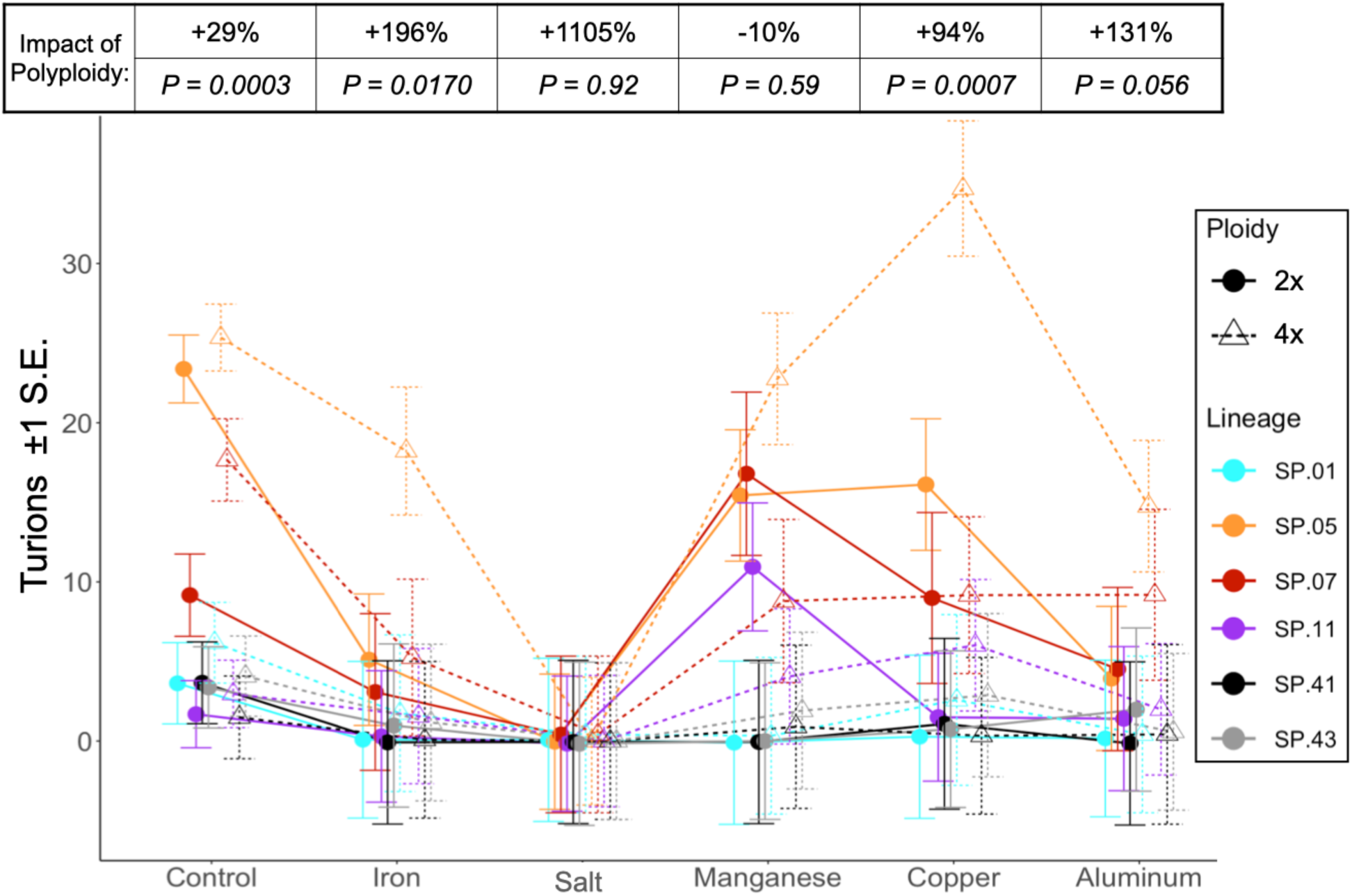
Total turion production (estimated marginal means ± 1 S.E.) separated by ploidy and genetic lineage across the six pollutant treatments (including the control). The table above shows the % difference of neopolyploid (4x) relative to diploid (2x) and P values from the six planned contrasts between diploids and neotetraploids for each pollutant treatment. Point positions were dodged for clarity.

### Neotetraploids often are more tolerant to pollutant stress

When tolerance is estimated as population growth under pollutant stress divided by that under control conditions, neopolyploidy generally increased tolerance although this was also influenced by lineage (Fig. 3). Again, interactions between ploidy, lineage, and pollutant type were significant (Table S4). A planned contrast averaging across all pollutants and lineages found that polyploids are more tolerant (estimated marginal mean 0.678) than diploids (0.520, P < 0.0001). This positive impact of neopolyploidy is statistically supported for all pollutants (planned contrasts; +6% to +66%, Fig. 3) except for iron. Yet, genetic lineage had a strong impact, with neopolyploids in three lineages (SP.01, SP.05, and SP.11) having higher tolerance than their respective diploids under all stressors, neopolyploids in two lineages (SP.41 and SP.43) had rank order changes across pollutants, whereas neotetraploid SP.07 was uniformly less tolerant.

**Figure 3:**
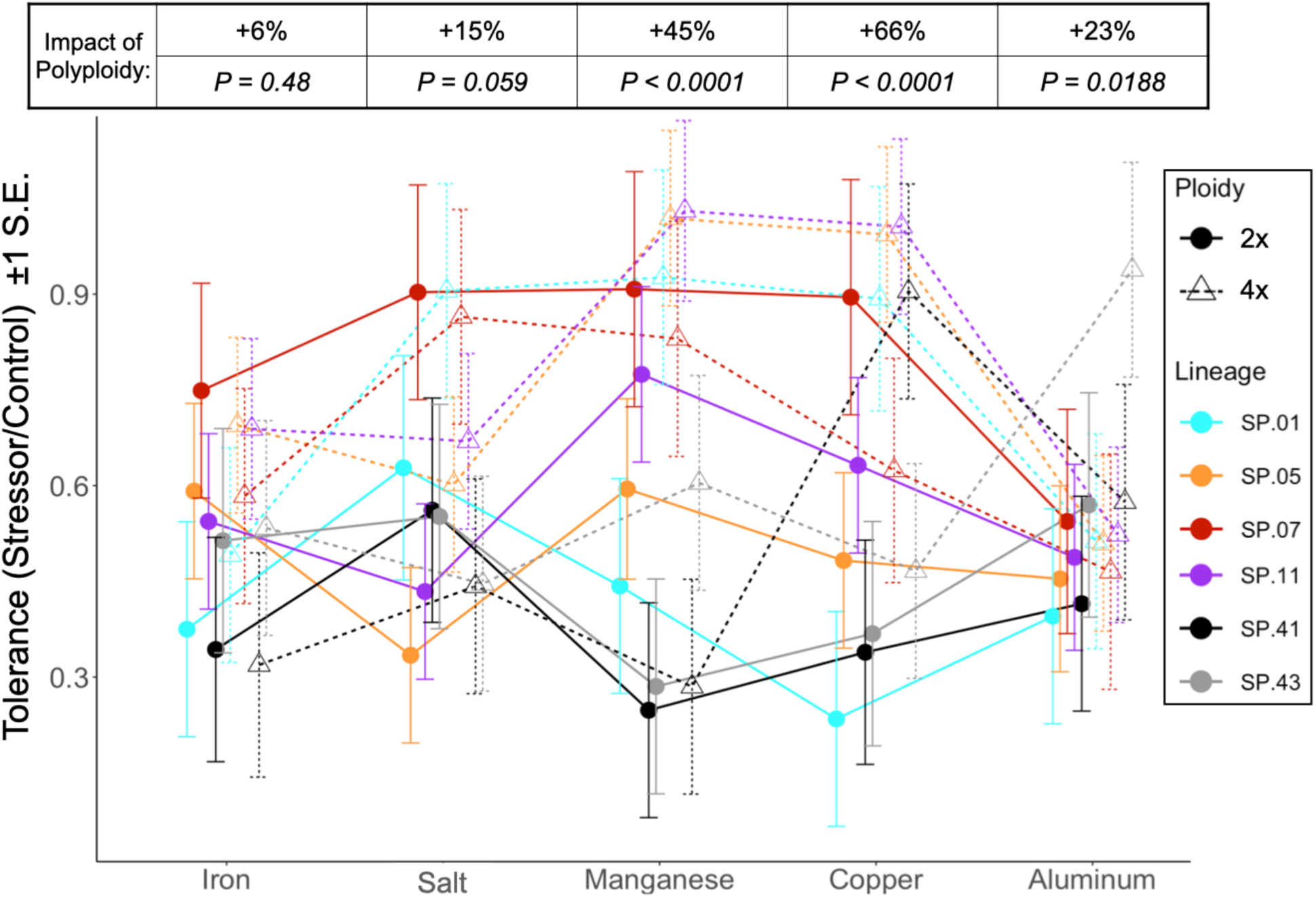
Tolerance of each pollutant (stressor/control, estimated marginal means ±1 S.E.) combining frond and turion growth across ploidy and genetic lineage. Tolerance values of 1 imply no impact of stressor on fitness. The table above shows the % difference of neopolyploid (4x) relative to diploid (2x) and P values from the five planned contrasts between diploids and neotetraploids for each pollutant. Point positions were dodged for clarity.

### Neotetraploids maintain fitness across pollutants better and showed fewer genetic tradeoffs in fitness under varied pollutants than their progenitor diploids

Polyploids maintained population growth across various stressors better than diploids as evidenced by both lower plasticity (measured as RDPI) and positive broad-sense genetic correlations. Ploidy, lineage and their interactions significantly impacted fitness plasticity across the five pollutants (Fig. 4; Table S5, all P < 0.0001). Overall, polyploids had lower fitness plasticity than diploids (0.268 vs 0.290) but this varied among lineages. In SP.01, SP.05, SP.11 and SP.41 neotetraploids maintained fitness better than their diploid progenitors. But the opposite was observed with SP.43. Finally, lineage SP.07 had the lowest and least variable fitness plasticity between ploidies (diploid vs neotetraploid: 0.190 and 0.193, Fig. 4).

**Figure 4:**
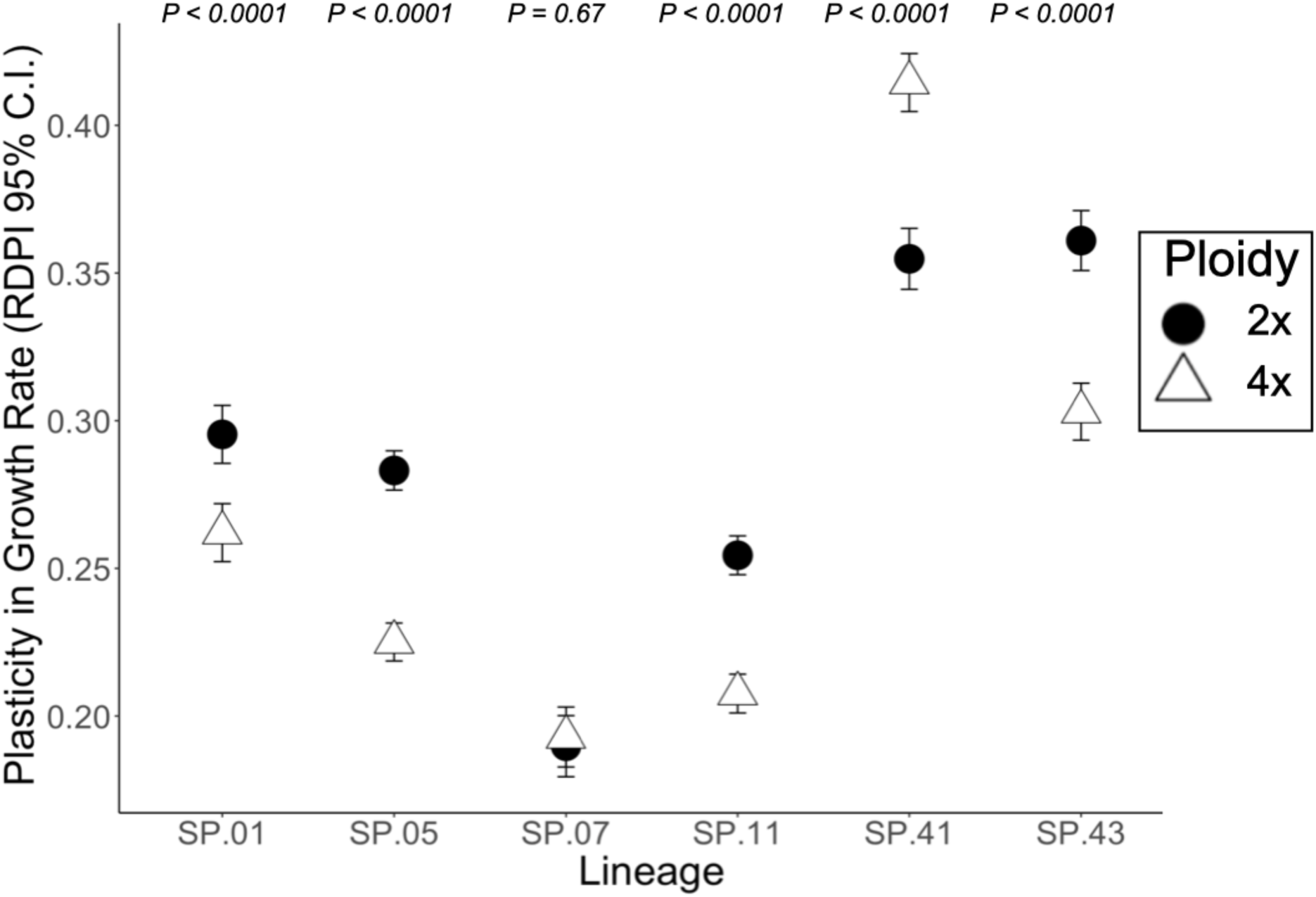
Plasticity in fitness across all pollutants (excluding the control) for each lineage and ploidy. Plasticity, quantified as the Relative Distance Plasticity Index (RDPI) of growth rates, combining fronds and turions, among the five pollutants. Values of 0 imply complete fitness maintenance (no plasticity) and 1 maximum plasticity. Post-hoc Tukey HSD test results are shown above each lineage pair.

Broad-sense genetic correlations in growth rate between pairs of metal pollutants were impacted by neopolyploidy. Correlations ranged from negative to positive in diploids but were all positive in neotetraploids (Fig. 5) but given the small sample size, only four individual correlations had P values between 0.001 and 0.07 (diploids: copper-salt −0.81, P = 0.05; iron-salt −0.77, P = 0.07; neotetraploids: aluminum-copper 0.96, P = 0.003; iron-manganese 0.92, P = 0.011). Nevertheless, across all correlations, a paired *t*-test revealed that diploids had significantly weaker and more negative correlations than polyploids (difference of −0.53, t = −3.09, df =9, P = 0.013).

**Figure 5:**
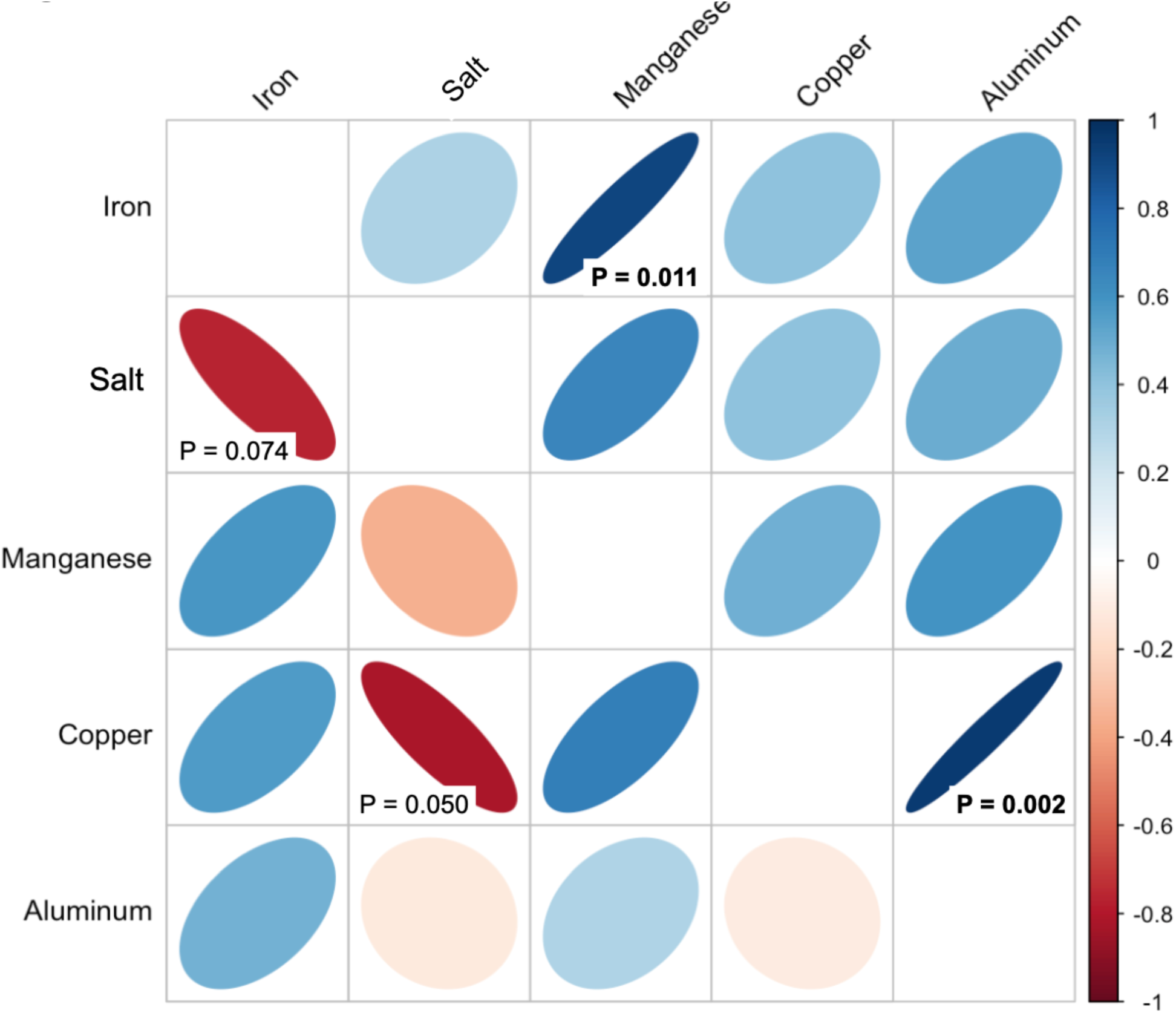
Broad-sense genetic correlations of population growth rate between pairs of pollutant environments (excluding the control). The lower triangle of the matrix includes Pearson’s correlations for diploids and the upper for tetraploids. P-values less than 0.075 are shown. Positive (negative) correlations are in blue (red) with darker shade reflecting strength of the correlation.

## DISCUSSION

Our exploration of the effects of neotetraploidy on duckweed population growth across five common urban pollutants provided support for two hypothesized advantages of polyploidy both in a general sense and with special regards to urban habitats. We found that relative to their diploid progenitors, neopolyploid duckweeds 1) are better stress tolerators and 2) show lower fitness plasticity across heterogenous pollutant environments and thus appear to have general purpose genotypes. And while the evidence for these generalizations is compelling, important complexities were also revealed. Specifically, ploidal responses varied by genetic lineage as well as by pollutant. Indeed, some stressful conditions reverse the ranking of ploidies. In the following paragraphs we explore in more depth the implications of both genetic- and environment-dependent advantages of neopolyploidy under urban pollution.

Our results demonstrated that stress conditions can upend fitness differences between the ploidies. Under favorable (control) conditions neotetraploid populations grew slower via fronds but produced more turions (Fig. 1 and 2). These results corroborate previous studies with synthetic neopolyploid duckweeds (Anneberg *et al*., 2023; Assour *et al*., *in Review*). But when exposed to pollutant stress, population growth rate of fronds in diploids was more similar to that of polyploids, however, the degree of change depended on pollutant and genetic lineage. Similar, three-way interactions dominated turion production. Interestingly, some pollutants suppressed a ploidal difference, manganese for fronds, iron and salt for turions. One contaminant even reversed the direction of diploid-neopolyploid difference: in copper polyploids performed significantly better than diploids in active propagule production. Similar, pollutant-dependent outcomes have been seen at the organismal level where salt and low nutrients and their combination lead to varying responses in two for *Arabidopsis* neotetraploids (Mattingly & Hovick, 2023). Much like our results, Bafort et al. (2023), using a different set of pairs of neotetraploid-diploid duckweed, found the advantage of polyploidy to be environment and genetic lineage specific. Neopolyploids showed an increase frond surface area at low concentration of salt. Yet their study did not measure turions. As we observed turion production, which is typically higher in neopolyploid duckweed, was suppressed by salt (Fig. 2), perhaps this gave rise to greater allocation by neopolyploids to fronds production under these conditions, and lower ploidal difference here as well (Fig. 1). Taken together these studies suggest that there are stressful conditions associated with urban activities that may allow neopolyploids to establish and persist locally. Whether neopolyploids are more common in urban environments remain unknown (Van Drunen & Johnson, 2022). Indeed, because neopolyploid duckweeds invest more heavily in allocation to future propagules in duckweeds (turion production) across most pollutant conditions tested, neopolyploids may be even more likely to persist in urban environments with severe winters (Anneberg *et al*., 2023).

Consistent with these fitness results, we also found that neopolyploidy leads to higher tolerance across most pollutants and genetic lineages. Relative to control conditions, polyploids are better able to maintain their total population growth rate (fronds and turions; Fig. 3). Although this result is influenced by the fact that diploids have higher fitness in control conditions (Fig. 1), it may indicate that polyploids are better able to handle stress but at a cost that is only apparent in ideal, uncontaminated growth conditions, which are increasingly rare in the Anthropocene. Whether neopolyploid’s higher tolerance is due to higher capacity for storage, greater detoxification, or other mechanisms remains to be determined with cellular morphometric and enzymatic studies, such as those reviewed in Tossi et al. (2022).

We found that relative to diploids, neopolyploids had lower plasticity in population growth rate across stress conditions indicating a maintenance of fitness across heterogeneous pollutants. While on first glance this may seem counter to other predictions and findings of increased phenotypic plasticity in polyploids (Parisod *et al*., 2010; Spoelhof *et al*., 2017; Van de Peer *et al*., 2017; Wei *et al*., 2019; Mattingly & Hovick, 2023), it is important to note that these other studies focus on plasticity in functional *traits* not *fitness* and indeed higher trait plasticity is expected to buffer fitness and thus lead to lower fitness plasticity (see Wei *et al*., 2019). Here we did not measure specific traits so we do not know how or what phenotypic changes might contribute to the fitness maintenance seen in polyploids. These could be changes in frond size, thickness, or photosynthetic activity seen in Bafort *et al*. (2023) in response to salt and cadmium pollution, or changes in the numerous other mechanism of detoxification of pollutants observed in diploid duckweeds (Ekperusi *et al*., 2019; Huber *et al*., 2021). Work to make these mechanism-function connections are a key next step.

Nevertheless, the shifts in broad-sense genetic correlations of fitness across pollutants from negative to positive (Fig. 5) for neopolyploids gives rise to the intriguing possibility that neopolyploidy leads to an instantaneous buffering from constraints. And while we acknowledge the low power of this test, these seem not to be driven by similar types of pollutants as the significant pairs were from different classes (alkaloids, heavy metals, and salt) rather than within one. Such an outcome could reflect a global advantage of larger cells (*gigas* effect; Doyle & Coate, 2019) and thus increased storage of toxic substances by neopolyploids. They could however also reflect upregulation or rewiring of shared antioxidant pathways (Lu *et al*., 2020), so we encourage greater exploration of this idea with increased number of genetic lineages, classes and concentrations of pollutants while still employing rigorous fitness measures. Integrated studies of stress tolerance across levels from cellular to organismal level will be crucial for understanding the polyploidy-stress relationship across systems (Wei *et al*., 2020).

Across our study we found a strong impact of the interaction between genetic lineage and ploidy within one or among different pollutants. The effect of ploidy on performance, tolerance, and fitness plasticity were all influenced by genetic lineage either quantitatively or sometimes qualitatively. Even though genomic studies suggest that *S. polyrhiza* has low genetic diversity (Ho *et al*., 2019; Xu *et al*., 2019), both our study with lineages of NE USA and that of Bafort et al. (2023) with lineages from different continents demonstrated pronounced difference among lineages in the effects of polyploidization. This disconnect suggests that functional differences within diploid *S. polyrhiza* may derive from epigenetic variation that can be altered by autopolyploidization along with genetic variation (Chen, 2007; Huber *et al*., 2021) and strongly influence stress tolerance and population growth. Regardless of the mechanism, our results join that of several others demonstrating the importance of genetic variation in diploid progenitors on neopolyploid morphology (Wei *et al*., 2020), population growth (Anneberg *et al*., 2023) and response to abiotic (Wei *et al*., 2020; Bafort *et al*., 2023) and biotic interactions (Forrester *et al*., 2020; Anneberg *et al*., *In Press*; Assour *et al*., *in Review*). And provide additional support for the idea that repeated evolution of polyploids may be key to their success (Soltis & Soltis, 1999; Kolář *et al*., 2017; Wei *et al*., 2020). Moreover, this type of variation is especially important in the context of eco-evolutionary dynamics that could occur in urban environments (Verrelli *et al*., 2022), because not only does it suggest that the impact of ploidy is modulated by the genetic background of the progenitor diploid but it also indicates that there is ample opportunity for natural selection. Finally, because we have shown here that the degree and direction of ploidal fitness difference varies among the pollutants means there is the potential for heterogeneity in ploidal dominance within urban environments depending on specific pollutants environment.

In conclusion, our study corroborates that of others suggesting that the combination of environmental perturbations and independent polyploidization events maybe crucial for the establishment of polyploids (Fawcett *et al*., 2009; Wei *et al*., 2020;

Anneberg *et al*., 2023). This may be even more important in the 21^st^ century as urban stressors may elevate both polyploid production and stress levels (Van Drunen & Johnson, 2022).

## AUTHOR CONTRIBUTIONS

MMT, TZ, NK, and KW created and deployed the curriculum, and helped collect/collate/clean the data. All authors contributed to the design of the experiment. MMT and TLA analyzed and interpreted the data and wrote the manuscript. All authors contributed to manuscript revisions.

## ACKNOWLEDGEMENTS

We would like to thank the students of Duckweed Survivor (BioSci 0057, Spring 2023 semester) and instructors (M. Amour, D. Bisi, S. Gess, N. Kanmanii, N. Kaufmann, A. Martin, A Nigam, K. Parks, B. Reed, J. Robertson, S. Stuckman, K. Wagner, and C. Wiesner) for conducting the experiment and K. Butela, K. Wozniak, and L. Rzodkiewicz for input on the course development, S. Bhattacharya, J. Robertson, and K. Kroeger for preparing duckweed populations for the course, E. O’Neill, J. Kerstetter, T. Anneberg, and A. Burr for help creating and maintaining the diploid-polyploid lineages. This work was supported by Dietrich School of Arts and Sciences, Department of Biological Sciences, NSF grant DEB-1935410 to MMT and DEB-2027604 to TLA, Mellon Fellowship to TZ.

## SUPPLEMENTAL MATERIALS

**Table S1:** Collection location of the six lineages as well as their microsatellite alleles values using loci from Xu *et al*. (2018) and Kerstetter *et al*. (2023). Complete or partial allele values of these lineages were previously published in Kerstetter *et al*. (2023) and or Anneberg *et al*. (2023).

| Lineage | Sampling location | GPS coordinates (Decimal Degrees) | Sp10 | Sp12 | Sp25 | Sp42 | Sp47 | Sp51 | Sp53 | SP.7286 | SP.7814 | SP.Pso31 |
| --- | --- | --- | --- | --- | --- | --- | --- | --- | --- | --- | --- | --- |
| **SP.01** | Beaver County, PA, USA | 40.7298, -80.3602 | 198/212 | 274/292 | 201/215 | 197/211 | 248/252 | 272/286 | 262/262 | 386/390 | 222/222 | 240/240 |
| **SP.05** | Allegheny County, PA, USA | 40.6210, -79.8268 | 182/182 | 270/270 | 209/223 | 185/189 | 256/290 | 278/278 | 264/268 | 390/390 | 222/226 | 246/255 |
| **SP.07** | Allegheny County, PA, USA | 40.6208, -79.8281 | 182/182 | 268/274 | 209/213 | 181/189 | 256/288 | 268/280 | 266/268 | 390/390 | 220/220 | 246/255 |
| **SP.11** | Allegheny County, PA, USA | 40.6206, -79.8271 | 182/182 | 266/290 | 171/235 | 177/185 | 284/288 | 280/306 | 268/280 | 390/390 | 224/224 | 246/255 |
| **SP.41** | Butler County, PA, USA | 40.9716, -80.0186 | 208/212 | 290/290 | 201/219 | 197/209 | 252/252 | 270/290 | 262/268 | 388/392 | 222/222 | 243/243 |
| **SP.43** | Andover Township, Ohio, USA | 41.6064, -80.5401 | 208/214 | 290/298 | 199/199 | 187/195 | 266/266 | 270/284 | 262/262 | 390/390 | 222/222 | 240/240 |

**Table S2:** Statistical results of best fitting model testing the impact of ploidy, lineage, and pollutant treatment (including the control) on the population growth rate of fronds.

| Factor | numDF | denDF | F-value | P-value |
| --- | --- | --- | --- | --- |
| Intercept | 1 | 3201 | 2796.5412 | <0.0001 |
| Ploidy | 1 | 3201 | 371.6237 | <0.0001 |
| Lineage | 5 | 3201 | 93.5696 | <0.0001 |
| Pollutant | 5 | 3201 | 166.7282 | <0.0001 |
| Ploidy:Lineage | 5 | 3201 | 36.3606 | <0.0001 |
| Ploidy:Pollutant | 5 | 3201 | 55.5983 | <0.0001 |
| Lineage:Pollutant | 25 | 3201 | 21.2083 | <0.0001 |
| Ploidy:Lineage:Pollutant | 25 | 3201 | 8.9361 | <0.0001 |

**Table S3:** Statistical results of best fitting model testing the impact of ploidy, lineage, and pollutant treatment (including the control) on turion production.

| Factor | numDF | denDF | F-value | P-value |
| --- | --- | --- | --- | --- |
| **Intercept** | 1 | 1467 | 8.26829 | **0.0041** |
| Ploidy | 1 | 1467 | 2.89271 | 0.089 |
| **Lineage** | 5 | 1467 | 71.93508 | **<0.0001** |
| Pollutant | 5 | 1467 | 1.09236 | 0.362 |
| **Ploidy:Lineage** | 5 | 1467 | 5.30969 | **0.0001** |
| Ploidy:Pollutant | 5 | 1467 | 0.16797 | 0.974 |
| **Lineage:Pollutant** | 25 | 1467 | 5.92899 | **<0.0001** |
| **Ploidy:Lineage:Pollutant** | 25 | 1467 | 2.60562 | **<0.0001** |

**Table S4:**
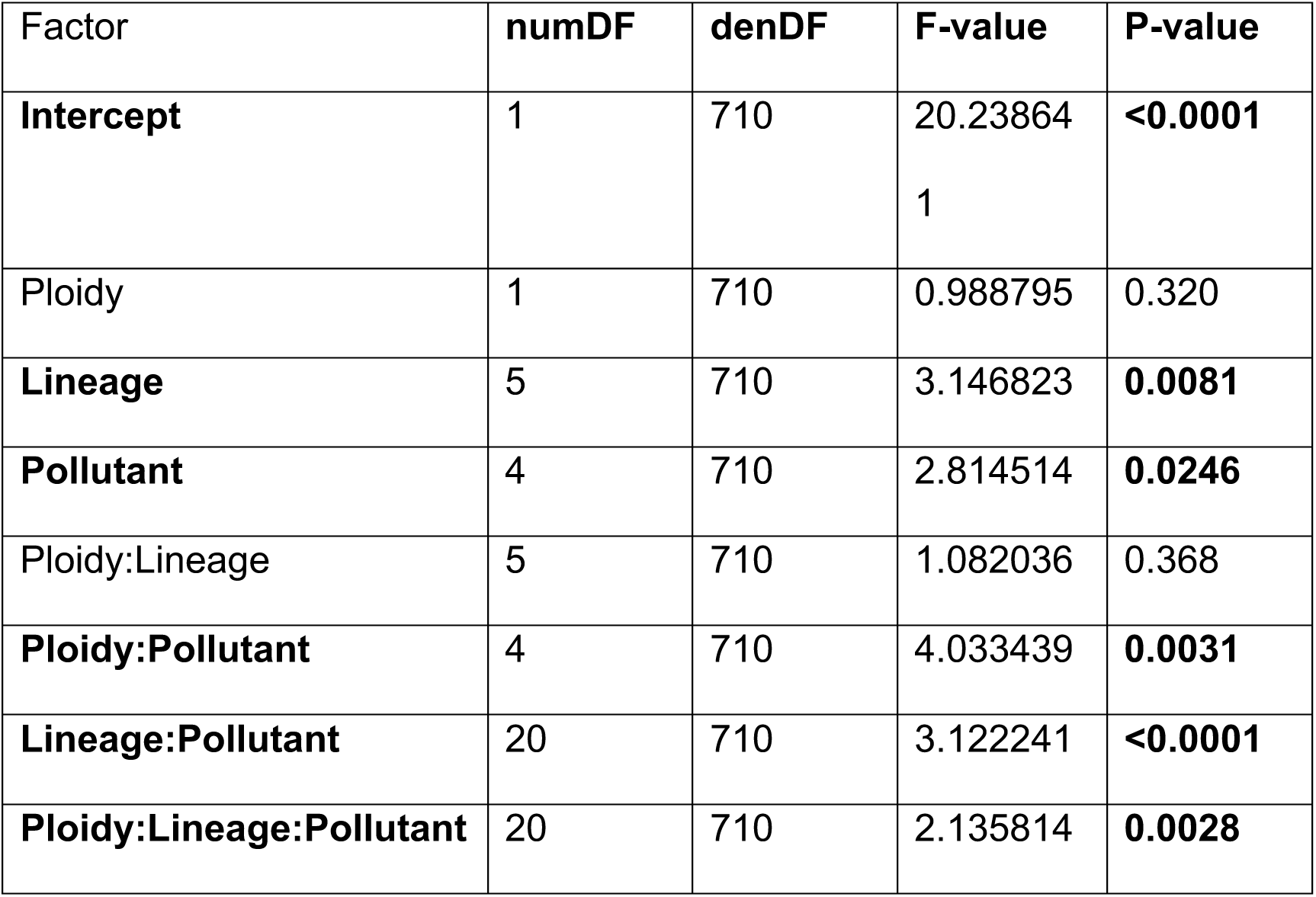
Statistical results of testing the impact of ploidy, lineage, and pollutant on tolerance across fronds and turions combined.

| Factor | numDF | denDF | F-value | P-value |
| --- | --- | --- | --- | --- |
| **Intercept** | 1 | 710 | 20.23864<br>1 | **<0.0001** |
| Ploidy | 1 | 710 | 0.988795 | 0.320 |
| **Lineage** | 5 | 710 | 3.146823 | **0.0081** |
| **Pollutant** | 4 | 710 | 2.814514 | **0.0246** |
| Ploidy:Lineage | 5 | 710 | 1.082036 | 0.368 |
| **Ploidy:Pollutant** | 4 | 710 | 4.033439 | **0.0031** |
| **Lineage:Pollutant** | 20 | 710 | 3.122241 | **<0.0001** |
| **Ploidy:Lineage:Pollutant** | 20 | 710 | 2.135814 | **0.0028** |

**Table S5:** Analyses of plasticity in growth rate (combining fronds and turions) across the five pollutants (not the control) for each lineage calculated as an RDPI. First table presents results from the linear model. The second table shows estimated marginal means of RDPI values for each sublineage growing in the metal stressors. Values near 0 imply fitness equivalence (no plasticity) and 1 maximum plasticity.

| Factor | DF | Mean Sq | F-value | P-value |
| --- | --- | --- | --- | --- |
| Ploidy | 1 | 152.775 | 152.775 | <0.0001 |
| Lineage | 5 | 421.194 | 421.194 | <0.0001 |
| Ploidy:Lineage | 5 | 47.022 | 47.022 | <0.0001 |

*Post-Hoc test*
| Lineage | Ploidy | Mean | S.D. | S.E. | Tukey post-hoc groups |
| --- | --- | --- | --- | --- | --- |
| SP.01 | 2x | 0.295 | 0.184 | 0.00493 | cd |
| SP.01 | 4x | 0.262 | 0.162 | 0.00434 | e |
| SP.05 | 2x | 0.283 | 0.204 | 0.00371 | d |
| SP.05 | 4x | 0.225 | 0.163 | 0.00287 | f |
| SP.07 | 2x | 0.19 | 0.127 | 0.0036 | g |
| SP.07 | 4x | 0.193 | 0.133 | 0.00369 | g |
| SP.11 | 2x | 0.254 | 0.209 | 0.00376 | e |
| SP.11 | 4x | 0.208 | 0.136 | 0.00245 | g |
| SP.41 | 2x | 0.355 | 0.236 | 0.00666 | b |
| SP.41 | 4x | 0.414 | 0.24 | 0.00644 | a |
| SP.43 | 2x | 0.361 | 0.227 | 0.0063 | b |
| SP.43 | 4x | 0.303 | 0.202 | 0.00532 | c |

**Figure S1:**
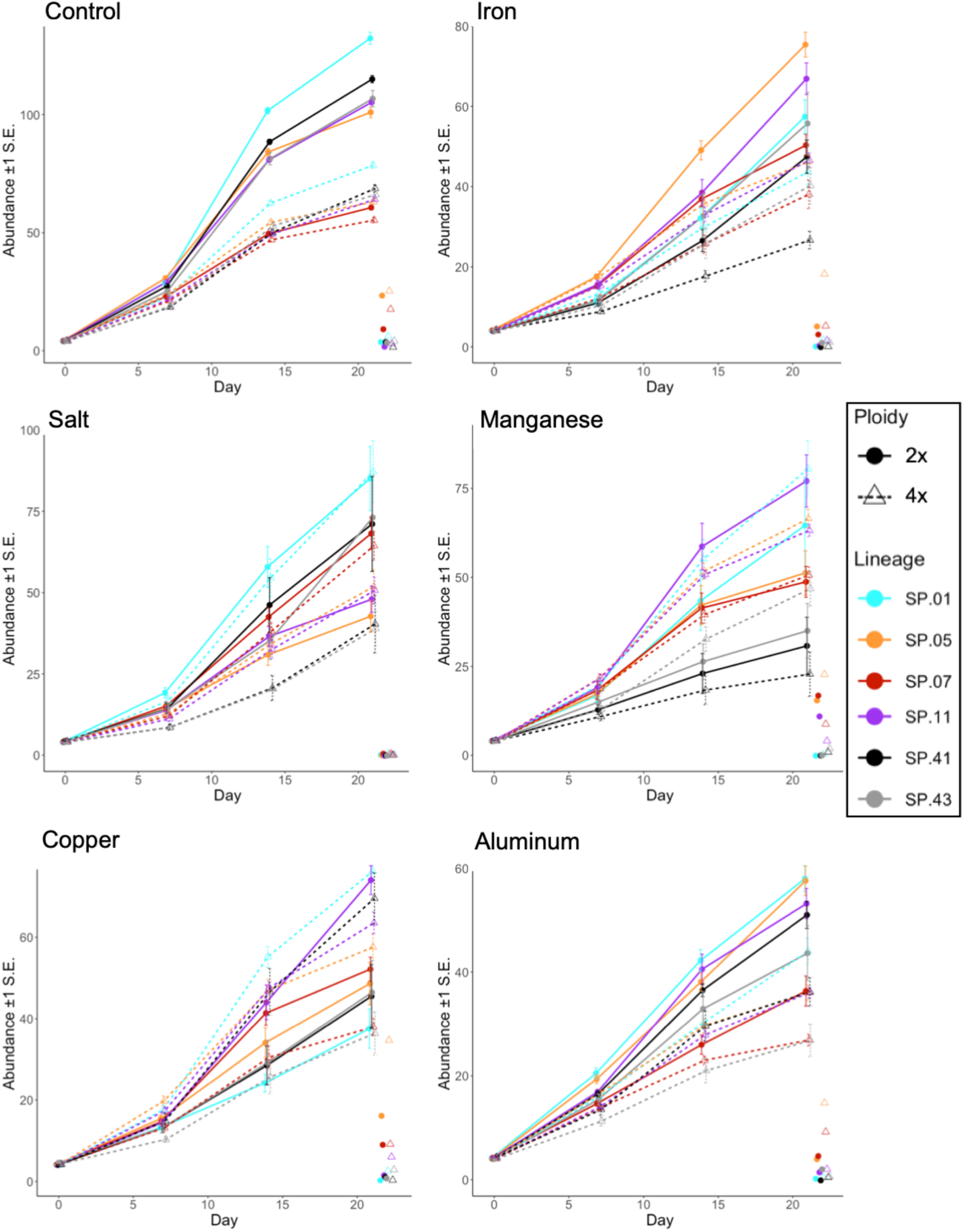
Time series of the abundance of fronds over time (estimated marginal means ±1 S.E.) in each pollutant treatment separately. In addition, the total number of turions are illustrated as a separate point without error bars for clarity.

